# Engineered Cardiac Microtissue Biomanufacturing Using Human Induced Pluripotent Stem Cell Derived Epicardial Cells

**DOI:** 10.1101/2024.05.13.593960

**Authors:** Kirk Butler, Saif Ahmed, Justin Jablonski, Tracy A. Hookway

## Abstract

Epicardial cells are a crucial component in constructing *in vitro* 3D tissue models of the human heart, contributing to the ECM environment and the resident mesenchymal cell population. Studying the human epicardium and its development from the proepicardial organ is difficult, but induced pluripotent stem cells can provide a source of human epicardial cells for developmental modeling and for biomanufacturing heterotypic cardiac tissues. This study shows that a robust population of epicardial cells (approx. 87.7% WT1^+^) can be obtained by small molecule modulation of the Wnt signaling pathway. The population maintains WT1 expression and characteristic epithelial morphology over successive passaging, but increases in size and decreases in cell number, suggesting a limit to their expandability in vitro. Further, low passage number epicardial cells formed into more robust 3D microtissues compared to their higher passage counterparts, suggesting that the ideal time frame for use of these epicardial cells for tissue engineering and modeling purposes is early on in their differentiated state. Additionally, the differentiated epicardial cells displayed two distinct morphologic sub populations with a subset of larger, more migratory cells which led expansion of the epicardial cells across various extracellular matrix environments. When incorporated into a mixed 3D co-culture with cardiomyocytes, epicardial cells promoted greater remodeling and migration without impairing cardiomyocyte function. This study provides an important characterization of stem cell-derived epicardial cells, identifying key characteristics that influence their ability to fabricate consistent engineered cardiac tissues.

## Introduction

Cardiovascular disease remains the leading cause of death worldwide, in part because of the heart’s limited capacity for self-repair. In the event of a heart injury, such as myocardial infarction, the myocardium responds by replacing the damaged region with scar tissue composed of extracellular matrix, primarily collagen [1]. This scar tissue allows the heart to maintain its structural integrity, preventing catastrophic heart failure in the short term, but permanently weakening the heart. The extracellular matrix of the scar is mostly secreted by activated myofibroblasts [2]. Myofibroblasts are derived from a population of resident fibroblasts in the heart, approximately 80% of which are epicardial in origin[2], [3], highlighting the importance of the epicardium in heart repair.

The epicardium is the outermost layer of the heart wall, located superficial to the myocardium. During development, the epicardium forms from the more primitive proepicardial organ (PEO) which grows alongside the myocardium until the two structures come into contact. In humans, leading cells from the PEO proliferate across the surface of the myocardium in a monolayer between Carnegie stages 12-14 until the heart is covered [4–7]. Studies of human epicardial cell migration have been rare, leaving animal models to shed light on these cells’ behavior. In explanted zebrafish and mouse tissue, epicardial cells expand over a surface as a contiguous sheet, migrating in a stratified pattern: a wave characterized by large, multinucleated cells is followed by a wave of predominantly small, mononucleated cells [8]. This migration pattern allows epicardial cells to rapidly cover the surface of the myocardium, allowing the epicardium to recover from injury and contribute to repair of the damaged tissue.

Epicardial cells are mesothelial in character, forming a monolayer of cells joined together by abundant tight junctions [7, 9]. Epicardial cells and their differentiated progeny—collectively referred to as epicardium-derived cells—can be identified by expression of the transcription factor Wilm’s Tumor 1 (WT1). WT1 acts as a central regulator to many key functions of the epicardium; during development it is necessary for vascularization of the heart [10, 11], and its upregulation following myocardial infarction is associated with ECM construction, epicardial-mesenchymal transition, and proliferation [11]. Epicardial cells are proliferative, migratory epithelial cells capable of epithelial-mesenchymal transition and ECM modulation which play a major role in proper cardiac development and repair after injury. Epicardial cells can transition into cardiac fibroblasts and smooth muscle cells which migrate into the myocardium and contribute to its structure and repair by deposition of ECM proteins, notably collagens, fibronectin, and laminin.

Because of their anatomical location and developmental origin, study of human epicardial cells is difficult. Induced pluripotent stem cells (iPSC) provide a means to study the behavior and interactions of human cells *in vitro*. Over the past decade, advances in iPSC differentiation have enabled researchers to generate robust populations of many different cell types. Recent studies have produced reliable protocols for stem cell differentiation to a variety of heart cell types, including ventricular cardiomyocytes [12], [13–18], atrial cardiomyocytes [16,19], epicardial cells [9, 20–22], cardiac endothelial cells [23, 24], cardiac fibroblasts [25–27], and nodal cells. [28–30] These iPSC-derived cells can be used to model human heart cell behavior and wound repair processes. Spherical 3D engineered heart microtissues (MT) have been shown to improve upon typical 2D cell culture in terms of functional maturity of cardiomyocytes [31–34], making them a more realistic model system. Additional supporting non-myocytes can improve on this approach by reproducing a more realistic mixture of the cell types found within the heart [34–36]. Recent studies have highlighted the importance of epicardial cells within engineered heart models with respect to their secretion of beneficial cytokines [37] or their role ischemic disease and recovery [21, 38, 39].

This study systematically evaluates the behavior of iPSC-derived epicardial cells (hereafter referred to as EpiC) in 2D culture as well as within 3D engineered heart tissue constructs, including heterotypic co-culture with iPSC-derived cardiomyocytes. The purpose of the study is to identify critical culture parameters under which EpiC can be most readily expanded and best utilized for the fabrication of engineered cardiac tissues and to increase our understanding of the role of epicardial cells in development and wound repair, both independently and in context with cardiomyocytes. The spherical EpiC microtissue model can be used to identify novel behaviors of human EpiC as well as to model the proepicardial organ and migration of EpiC, which would otherwise prove highly challenging.

## Methods

### Pluripotent stem cell culture

WTC11 human induced pluripotent stem cells modified with GCaMP6f (Gladstone Institutes) or mEGFP-conjugated TJP1 (Coriell Institute) were cultured as previously described [40]. Cells were maintained (37 °C, 5% CO_2_) in Essential 8 media (Gibco) on 23 µg/cm^2^ Matrigel (Corning) coated dishes. At 70% confluence, hiPSC were singularized using Accutase (Innovative Cell Technologies), and seeded at 14,000 cells/cm^2^. During seeding, the culture media was supplemented with 10 μM ROCK inhibitor (Y-27632, Selleckchem).

### Stem cell differentiation

Human iPSCs were seeded at 36,000 cells/cm^2^ and differentiated to epicardial cells (EpiC) following a modified Wnt modulation protocol [20]. At 85% confluence, the media was changed to RPMI with B27 supplement minus insulin (Gibco), 12 μM GSK3 inhibitor (CHIR99021, Selleckchem). CHIR99021 was removed after 24 hours (D1). At D3, 5 μM Wnt inhibitor (IWP2, Selleckchem) was added for 48 hours and removed on D5. On D6, cells were singularized with Accumax (Innovative Cell Technologies) and cryopreserved at 2x10^6^ cells/mL in media containing 30% FBS (Sigma-Aldrich) and 10% DMSO (Sigma-Aldrich) with 10 μM Y-27632.

Upon thaw, cells were seeded at 25,000 cells/cm^2^ onto gelatin-coated polystyrene (10 µg/cm^2^, Millipore). Epicardial media was formulated from Advanced DMEM/F-12 (Gibco), 1.3% GlutaMAX (Gibco), and 96 μg/mL L-Ascorbic Acid (Sigma-Aldrich) [20]. On D7 and D8, 3 μM CHIR99021 was added to the epicardial media and then removed again on D9 and D10. At D11, cells reached the epicardial phenotype and were used for experimentation. For further expansion, D11 EpiC were seeded onto gelatin-coated polystyrene at 25,000 cells/cm^2^ and cultured in epicardial media with 5 μM ALK inhibitor (A83-01, Sigma-Aldrich) to prevent spontaneous differentiation. At this point EpiC are referred to by passage number, not day of differentiation.

To produce cardiomyocytes, we followed a standard Wnt modulation protocol [12], identical to the above until D5. At D7, the media was changed to RPMI with B27 supplement (Gibco) and refreshed every two days. By D9, the cells contracted spontaneously.

### Immunofluorescent staining

For sequential study of EpiCs over long term maintenance, cells were seeded onto coverslips at each passage, and were then fixed using 4% paraformaldehyde. Cells were then permeabalized (0.1% TritonX-100 in PBS, 15 minutes), blocked (1.5% Normal Donkey Serum, 1 hr), exposed to primary and secondary antibodies (WT and ZO-1, see Supplementarty Table 1), secondary antibodies and counterstained with Hoechst (15 minutes). The samples were then imaged using Nikon Ti2.

### Flow cytometry

Samples of at least 10^6^ cells were singularized at D0 and D4 using Accutase and at D11 through Passage 6 using Accumax. D11 samples were acquired from both EpiC and cardiomyocyte differentiations from the same original D0 population. The samples were fixed using 4% paraformaldehyde for 15 minutes. For flow cytometry staining, working buffer (3 mg/mL BSA (Sigma-Aldrich) in DPBS with 0.01% Tween-20) was supplemented with 0.5% Triton X-100 (Sigma-Aldrich) for permeabilization and 1% Normal Donkey Serum (Sigma-Aldrich) for blocking. Three biologically independent samples of each condition were incubated for one hour using primary antibodies against Oct4, cardiac troponin T, and WT1 (all Abcam) followed by 45 minutes incubation with species-appropriate Alexa Fluor Plus 488 secondary antibodies (Invitrogen). A full list of antibodies and working dilutions can be found in Supplementary Table 1. All samples were run through a Bio-Rad ZE-5 flow cytometer for analysis with a minimum of 10,000 cells. The populations were gated using FlowJo software and the mean percentage of the three populations positive for each marker was reported. All percentages derived from flow cytometry results were reported as mean ± SEM.

### EpiC microtissue formation

Both D11 and P1 EpiCs were singularized by Accumax for 6 minutes and resuspended in epicardial medium with 1% FBS, 5 μM A83-01, and 10 μM Y-27632. The cells were seeded into an ultra low attachment round-bottom 96-well plate at a density of 10,000 cells/well. Plates was centrifuged at 200xg for 3 minutes then cultured for 7 days with 50% media volume exchanges every two days. The microtissues were imaged daily using phase microscopy (Nikon Ti-2). At D7 the tissues were fixed in 10% formalin for 30 minutes, embedded in HistoGel (Epredia), and processed for histology and immunohistochemistry. Five μm sections were stained with Harris Hematoxylin and Eosin Y (both Sigma-Aldrich), and immunostained for WT1 with a Hoechst (Invitrogen) counterstain.

### EpiC microtissue expansion and morphometrics

To assess EpiC migration, microtissues were assembled from D11 EpiC using the mEGFP-TJP1 tagged cell line. After 24 hours of formation, microtissues were transferred to flat-bottom 48 well plates coated with different extracellular matrix coatings (tissue culture polystyrene (TCPS), 10µg/cm^2^ gelatin, 23µg/cm^2^ Growth Factor Reduced Matrigel (Corning), and 2, 5, and 10μg/cm^2^ fibronectin (Corning), laminin (Corning), and collagen I (Corning) (product information found in Supplementary Table 2). After 24 hours, media was replaced with epicardial media containing 5 μM A83-01. Samples were imaged at 8, 16, 24, 48, and 72 hours.

To examine the morphology of migrating EpiC in more detail, D11 microtissues were expanded for 3 days 10 µg/cm^2^ gelatin as described above. The wells were fixed using 4% paraformaldehyde, permeabilized using Triton X-100 (Sigma-Aldrich), blocked in 1.5% Donkey Serum, and exposed to primary antibodies for WT1 (Abcam) and ZO1 (Invitrogen). A complete list of antibodies can be found in Supplementary Table 1. For characterization of expansion skirt area, the cells were stained using Alexa Fluor 488 conjugated Phalloidin (Invitrogen) and Hoechst. The stained cells were imaged using a Nikon Ti2 Eclipse fluorescence microscope with an Andor Zyla camera.

The outgrowth expansion region, or *skirt*, of each sample, was reconstructed using overlapping fields of view of either phase microscopy or immunofluorescence adjusted for maximum contrast and manually aligned (Adobe Photoshop). The reconstructed skirt areas were measured using the lasso tool in Image J. This same measurement was used to compute circularity of the skirts. The reconstructed skirt images from the ZO1 immunofluorescence were processed using J-SEG, a custom MATLAB script implementing the Cellsegm method [41]. Data produced from this software were used to characterize EpiC morphology (n=3 samples). This method was also used to determine individual cell size and examine Leader and Follower cell position. The overall range of distance from the skirt edge within the population was computed and a dividing line at half that distance was placed. The cells closer to the microtissue area were labeled as “Follower” EpiC and those closer to the skirt edge as “Leader” EpiC. The segmentation was repeated for composited images from each of three independent microtissue expansions on 10 µg/cm^2^ gelatin-treated surfaces, dividing each population at half its overall range of distance from the skirt edge. Binning the cells in this manner produced a consistent difference in proportion of large cells to small between the Leader and Follower subpopulations. The mean cell area of each sample’s Leader and Follower subpopulations was calculated, and the mean of these means was plotted.

### Formation of bulk spherical microtissues

Spherical EpiC microtissues were formed using agarose molds as previously described [40, 42]. Agarose molds were cast from inverse PDMS templates, then placed into 24 well plates along with 1mL of epicardial media. The plates were centrifuged at 2000xg and stored at 37°C until seeding.

D11 EpiC (GCaMP6f) were singularized using Accumax and resuspended in epicardial media with 1% FBS, 5 μM A83-01, and 10 μM Y-27632. CM were singularized at D15 Trypsin-EDTA. Cells were seeded into the agarose molds at 3,000 cells/well in RPMI with B27 supplement with 10 μM Y-27632. Microtissues were formed from cardiomyocytes alone (CMonly) or mixed at a ratio of 3:1 with epicardial cells (CMEpiC). Cells were centrifuged for 3 minutes at 200xg.

After 24 hours, microtissues were transferred into 100 mm non–tissue culture treated dishes containing 10 mL epicardial media with 5 μM A83-01. Dishes were maintained on an orbital rotary shaker at 60 rpm and media was replaced every 3 days. Phase images were taken daily over 7 days of culture. Calcium flux video recordings (100fps, 10-20 seconds) were obtained at D2, D4 and D7 of culture.

### Microtissue morphometrics

High-contrast bright field images of microtissues were imported into ImageJ and binarized using a threshold so that the resulting objects were subjected to a hole-fill method. Neighboring objects were manually segmented and the analyze particles function was used to measure cross sectional area, major and minor axes, and circularity.

### Contractile kinetics assessment of 3D microtissues

The time lapse calcium flux recordings of beating microtissues were imported into ImageJ. Within each recording, individual microtissues were isolated using a 100x100 pixel region of interest. The extracted recordings imported to the MUSCLEMOTION ImageJ plugin [43]. All settings were set to standard, specifying the 100 fps frame rate. After processing, the detected contraction profile and associated contraction characteristics for each microtissue were compared between groups.

### Statistical analysis

Statistical analysis of data was performed using GraphPad Prism 9. When multicolumn data were normally distributed and homoscedastic (as assessed by Shapiro-Wilk test and QQ plot for normality and Brown-Forsythe test or F-test for heteroscedasticity) differences in means were assessed by Welch’s t-test, one-way ANOVA with Dunnett’s post-hoc test, or two-way ANOVA with Tukey’s or Sidak’s post-hoc test, as appropriate. When data were normally distributed but heteroscedastic, we instead assessed differences in means by Brown-Forsythe test and Welch ANOVA with Dunnett’s T3 post-hoc test. When data were not normally distributed, we instead assessed differences in medians by Mann-Whitney test or Kruskal-Wallis test with Dunn’s post-hoc test. Trends were assessed using simple linear regression; we evaluated the resulting fit against the null hypothesis (*m* = 0) by F-test. For all statistical tests, *p-*values smaller than 0.05 were considered significant.

## Results

### Wnt modulation produces enriched epicardial populations

To evaluate efficiency of the differentiation protocols, iPSCs were directed toward either EpiC or CM (Figure 1A). Representative immunofluorescence images of characteristic markers for the EpiC and CM differentiation are shown in Figure 1B. Three days after seeding, at D0 of the differentiation, the iPSC population had strong expression of Oct4 around the edges of the colonies and a total of 85.2 ± 3.3% of the population was Oct4^+^ (Figure 1C). At this time there was negligible expression of cTnT (0.9 ± 0.1%) and WT1 (0.6 ± 0.1%), reflecting a population that is largely pluripotent, with some early differentiation initialized by high cell density at the beginning of EpiC induction. By D4, Oct4 expression was largely turned off (4.5 ± 0.7%) while expression of cTnT (1.8 ± 0.3%) and WT1 (0.4 ± 0.1%) remained minimal. Brachyury expression, a mesodermal marker, appeared transiently at this stage, indicating primitive streak induction. At D11 following the EpiC protocol, WT1 was significantly expressed (87.7 ± 0.2% WT1^+^) and the population contained low levels of cardiomyocytes (19.6 ± 1.4% cTnT^+^). Enrichment of the EpiC population compared to cardiomyocytes was observed at a ratio of 4.5:1 WT1:cTnT with negligible expression of Oct4 (1.2 ± 0.1%), Figure 1D. At D11 following the CM protocol, 62.1 ± 3.8% of the population stained positively for cTnT, with low expression of WT1 (8.8 ± 2.5%) and negligible expression of Oct4 (0.6 ± 0.1%). At D11 both EpiC and CM expressed Isl1, reflecting commitment to a cardiac lineage. Taken together, this data demonstrates an enrichment in epicardial phenotype following a directed differentiation.

**Figure 1.**
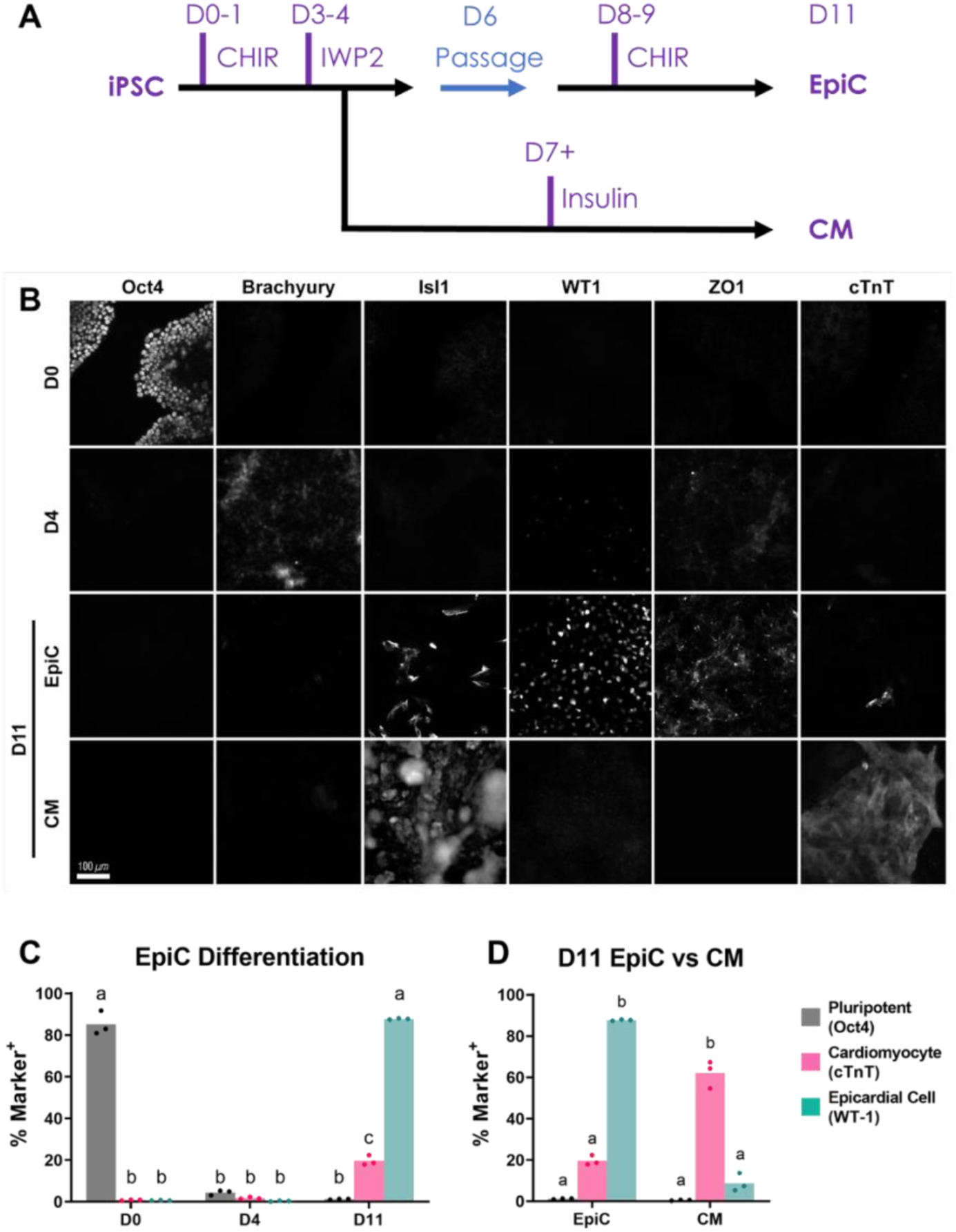
Differentiation of EpiC and CM populations. **(A)** Schematic of CM and EpiC differentiation protocols. **(B)** Representative images of marker expression throughout EpiC and CM differentiations (Scale = 100 µm). **(C)** Comparison of Oct4, cTnT, and WT-1 expression at D0, D4, and D11 of EpiC differentiation and **(D)** between the D11 EpiC and D11 CM populations. Different letters indicate statistically significant differences in means assessed by two-way ANOVA with Sidak’s post-hoc test.

### EpiC remain WT-1+ but decrease in cell density over prolonged culture

To evaluate the long-term viability of iPSC-EpiC for biomanufacturing purposes, EpiC were maintained in 2D culture through 5 passages. To observe changes in morphology and phenotype, WT1 and ZO1 immunocytochemistry and WT1 flow cytometry were performed on confluent EpiC just before each passage past D11 (Figure 2A). Cell count decreased at each successive passage, indicating a lack of cell growth, and by the fifth passage reseeding was no longer possible due to limited cell numbers. This downward trend was confirmed by linear regression (Figure 2B; R^2^ = 0.6909, *p* = 0.0002). Concurrently, the cells were found to increase in size at each passage, confirmed by linear regression (Figure 2C; R^2^ = 0.6115, *p* < 0.0001). This increase in size over time is consistent with the morphology observed across passages as shown in Figure 5A. Flow cytometry results over successive passages, however, indicated a maintenance of EpiC phenotype, are shown in Figure 2D. High WT1 expression was maintained over successive passages with mean expression between 70 and 90 percent of the population and analysis by simple linear regression found no statistically significant trend in expression (R^2^ = 0.2701, *p* = 0.2906). The high level of WT1 expression found in flow cytometry is consistent with WT1 immunofluorescence, and strong ZO1 expression further supports the characteristic mesothelial morphology of EpiC (Figure 2A). Taken together this shows that iPSC-derived EpiC maintain WT1 expression over prolonged culture, but increasing cell size limits expansion after D11.

**Figure 2.**
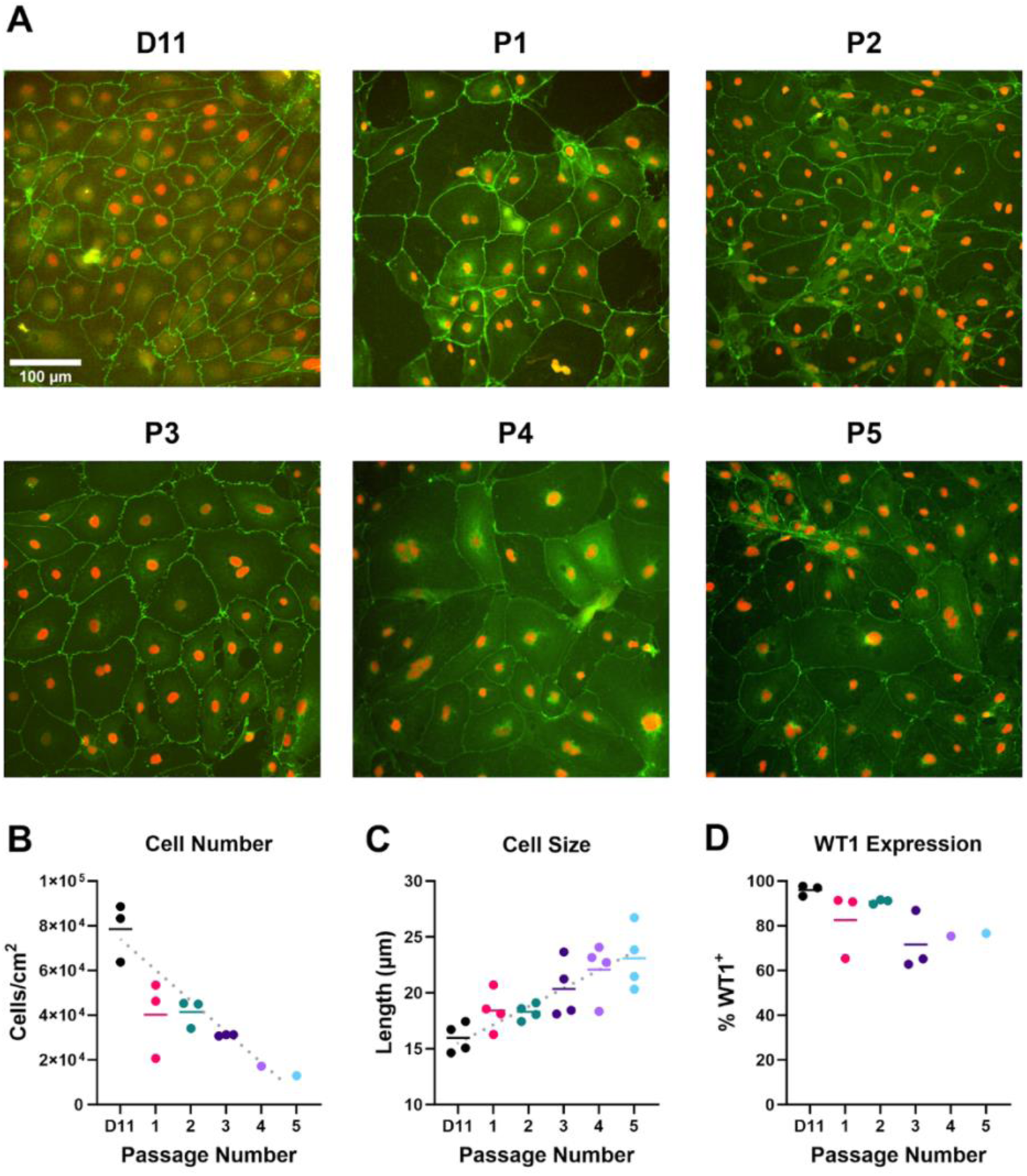
Phenotypic assessment of epicardial cells across multiple passages. **(A)** Representative WT-1 (red) and ZO-1 (green) staining of EpiC cells across 5 passages (Scale = 100 µm). **(B-D)** Quantitative assessment of cell density **(B)**, cell size **(C)**, and WT-1 expression **(D)** of EpiC cells over 5 passages. Linear regressions yielded no correlation.

### EpiC self-assemble into spherical 3D microtissues

Since both D11 and later-passage EpiC retain WT1 expression, both D11 and P1 EpiC were assembled into 3D microtissues to compare their performance in a model suitable for both biomanufacturing and studying the proepicardial organ. Both D11 and P1 EpiC successfully formed into spherical microtissues. After 24 hours of formation, microtissues from P1 EpiC were more loosely packed compared to microtissues from D11 EpiC, with an 88.3% greater major axis length. After 7 days of 3D culture, microtissues from D11 EpiC maintained their size whereas microtissues from P1 EpiC reduced in major axis length by 69.1% (Figure 3A). After the 7 days of 3D culture, the D11 microtissues were 78.3% larger than the P1 microtissues. Hematoxylin & Eosin staining revealed that microtissues from both D11 and P1 EpiC displayed irregular shapes, characterized by acellular spaces and a heterogeneous arrangement of cells. WT1 immunohistochemistry confirmed that most cells contained within all microtissues remained epicardial; however, stronger WT1 expression was observed in cells closer to the exterior of the microtissues assembled from D11 EpiC. (Figure 2B). The more consistent and ultimately larger size of D11 microtissues, coupled with a similar maintenance of phenotype and internal microtissue structure, suggests D11 EpiC are preferable for biomanufacturing and developmental modeling compared to P1 EpiC.

**Figure 3.**
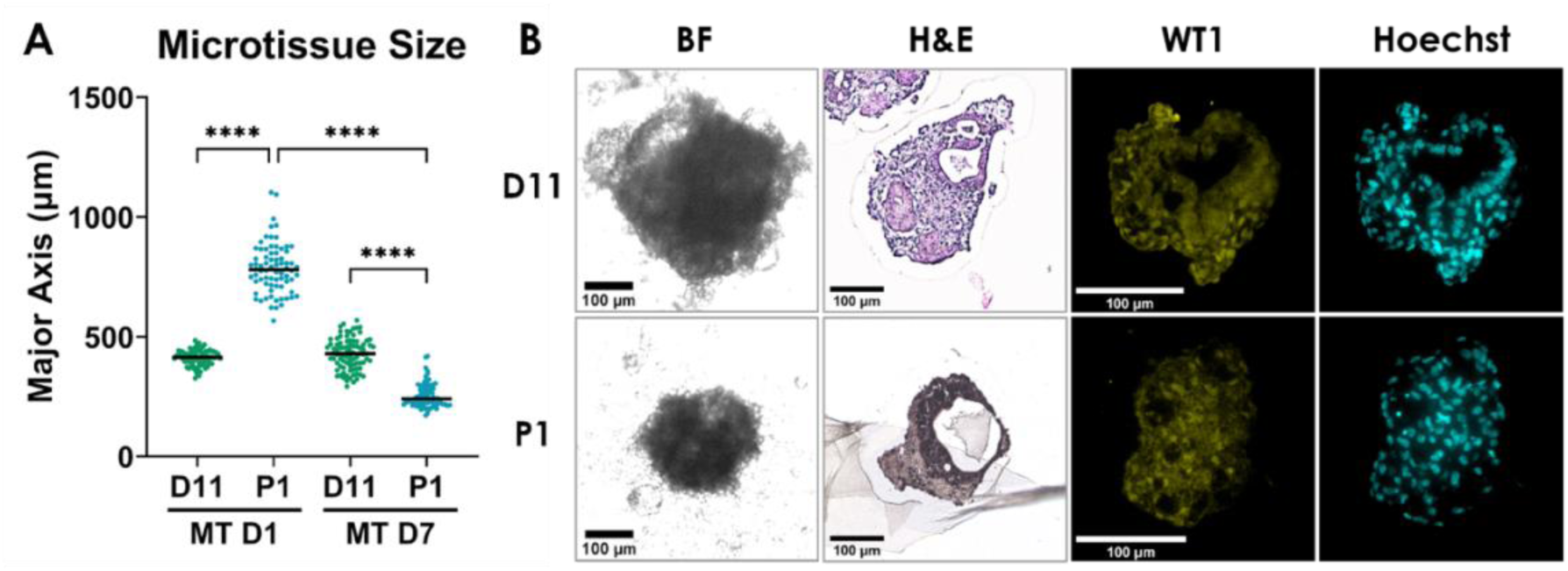
EpiC microtissue formation. **(A)** Size comparison (major axis length) of microtissues cultured for 1 and 7 days assembled from D11 or P1 EpiC. Asterisks indicate a statistically significant difference in mean assessed by Brown-Forsythe test and Welch ANOVA: (****) p < 0.0001 **(B)** Representative bright-field (BF), Hematoxylin & Eosin (H&E), WT1 immunohistochemistry (WT1), and Hoechst (Nuclei) images of microtissues seeded from D11 and P1 EpiC after 7 days of culture (Scale = 100 µm).

### Heart-relevant ECM surface treatments differentially impact EpiC migration

Having identified D11 EpiC as the most suitable population to form a 3D model of the proepicardial organ (PEO), the resulting tissues were used to evaluate the ability of EpiC to transition from a 3D culture environment into a 2D environment, modeling the outgrowth of cells from the PEO over the myocardium (Figure 4A). When deposited onto surfaces coated with extracellular matrix proteins, EpiC expanded outward from the microtissues in a migratory wavefront, forming a broadly circular “skirt” region composed of an epithelial-like EpiC monolayer. ECM coatings were selected based on their presence in cardiac ECM and use in propagation of cardiac cell types *in vitro* to simulate the ECM environment of the developing heart: gelatin (Gel), Matrigel (MG), collagen 1 (Col I), fibronectin (FN), and laminin (LN). The extent of EpiC migration after 3 days varied according to the ECM protein treatment with an initial migratory advantage observed in cells cultured on Col I, FN, and LN (Figure 4B, C). Untreated tissue culture polystyrene consistently yielded the smallest skirts. In most cases the difference in mean skirt area between conditions was not found to be statistically significant, though at 72 hours all three laminin-treated conditions improved significantly compared to TCPS (Figure 4D). All surface treatment types proved amenable to EpiC migration, consistent with the function of EpiC and their derivatives in remodeling the ECM environment of the heart.

**Figure 4.**
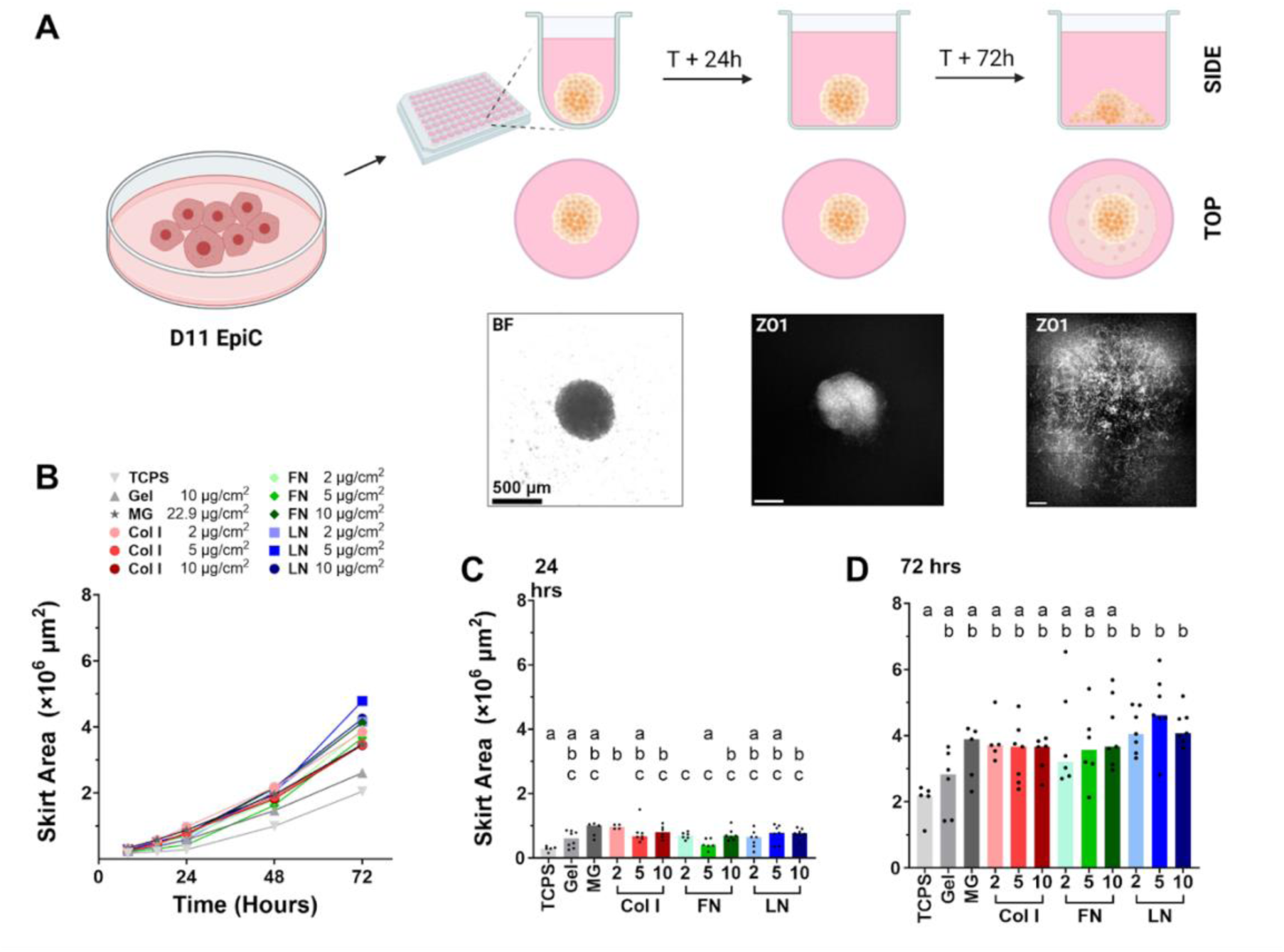
EpiC microtissue outgrowth on different ECM surfaces. **(A)** D11 EpiC were aggregated in round-bottom 96 well plates then transferred to flat surfaces for expansion. Images show brightfield or ZO1 fluorescence of corresponding stage of aggregation or outgrowth (Scale = 500 µm). **(B-D)** Measured skirt area over 72 hours of microtissue outgrowth on different extracellular matrix coatings (TCPS: tissue culture polystyrene, Gel: gelatin, MG: Matrigel, Col I: collagen I FN: fibronectin, LN: laminin). Comparison of skirt area over time **(B)** and in detail at 24 **(C)** and 72 hours **(D)**. Columns with no letters in common differ in mean (*p* < 0.05), as assessed by Brown-Forsythe and Welch ANOVA with Dunnett T3 post-hoc test.

**Figure 5.**
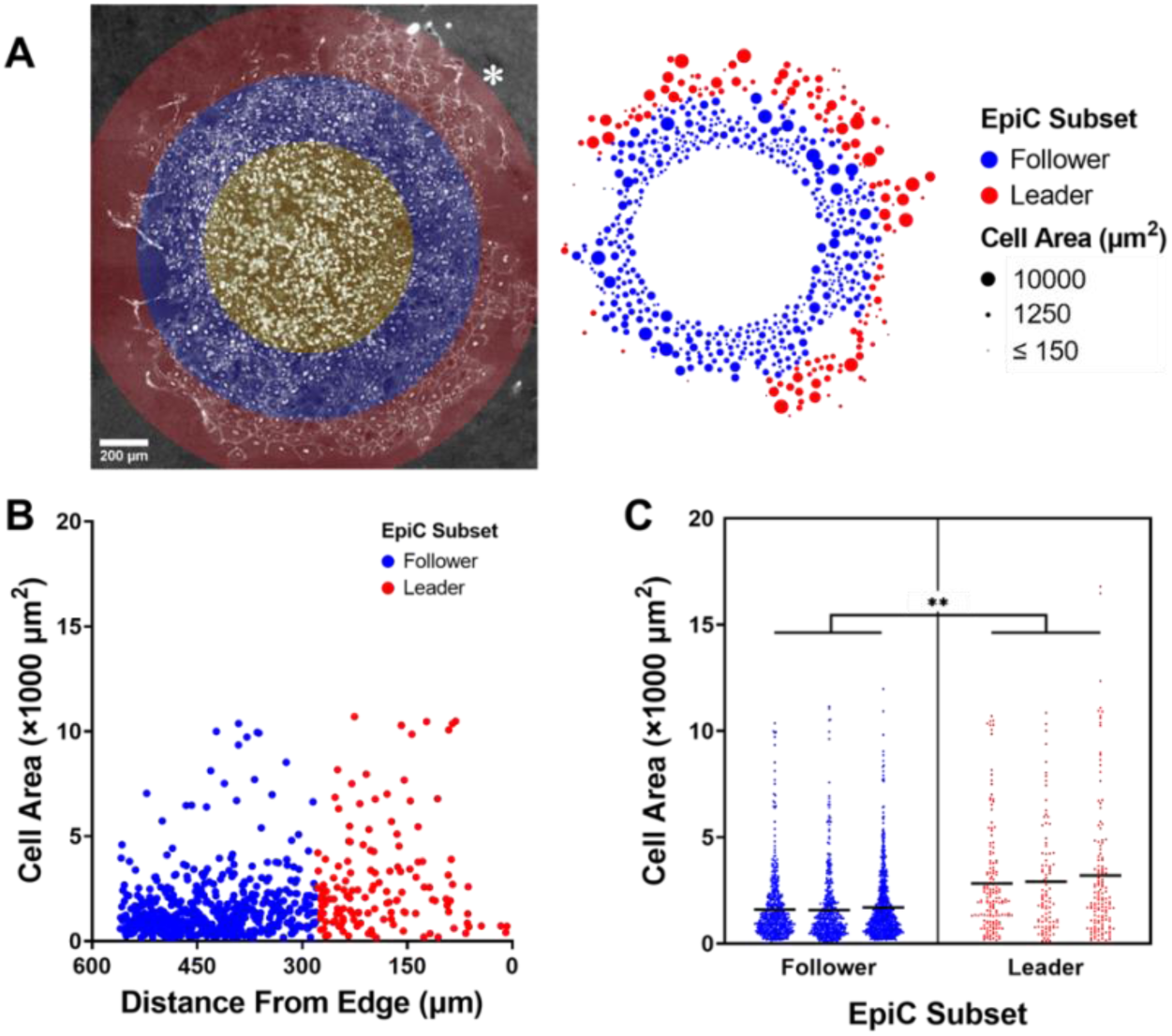
Populations of outgrowth cells. **(A)** Composited image (left) of expansion skirt for an EpiC microtissue after 7 days, showing cell borders (ZO1) and nuclei (WT1). Colored overlay indicates division of cells into Leader cells (red), Follower cells (blue), and microtissue region (yellow). Diagram (right) displays area of each cell (µm^2^). Color indicates division into Leader (red) and Follower (blue) populations. Scale = 200 µm **(B)** Scatter plot of cell population depicted in (A) indicating cell area (in µm^2^) as a function of distance from the edge of the expansion skirt (in µm). **(C)** Comparison of cell area (in µm^2^) between subpopulations across 3 independent samples. Horizontal bars represent subpopulation means and asterisks indicate a statistically significant difference in mean of means assessed by Welch’s t-test: (**) *p* < 0.005.

### EpiC migrate from 3D microtissues in a stratified wavefront to form a 2D monolayer

The skirt expanded from the central microtissue region to form an epithelial monolayer composed of varying sized EpiC cells. The EpiC population could be divided into two subpopulations, organized according to distance from the edge of the skirt (Figure 5A; skirt edge indicated by * in composite image). The cells closer to the leading edge of the skirt were termed “Leaders” and the cells trailing termed “Followers”. The overall mean cell areas of Leader and Follower cells were 2985 ± 111 µm^2^ and 1624 ± 36 µm^2^, respectively (mean ± SEM; assessed by Welch’s *t-*test with *p* = 0.0035 and *n* = 3). Because the range of cell area was large, we also examined the difference in overall median cell area of the two groups, and groups differed in this respect as well, with a median of 2043 µm^2^ for Leaders and 1148 µm^2^ for Followers. Across all three samples, 81.6% of cells were identified as Followers and the remaining 18.4% as Leaders. The preponderance of Follower cells reflects the higher cell density in the region closer to the microtissue as well as the difference in size between the groups. The difference in morphology suggests a subset of larger, more migratory EpiC within the population which lead the overall migration. This data indicate that a subpopulation of larger cells lead the wave front to migrate across a surface.

### Cardiomyocytes and EpiC form mixed heterotypic 3D microtissues

To evaluate the potential of EpiC as a component of an engineered heart tissue model, day 11 EpiC were incorporated in a 3:1 mixture of cardiomyocytes:EpiC in spherical 3D microtissues (CMEpiC) and compared with microtissues composed only of cardiomyocytes (CMonly). Both the mixed CMEpiC and the CMOnly populations successfully formed microtissues. By D4 of microtissue culture, CMEpiC were significantly larger than CMonly, despite containing the same initial number of cells (Figure 6C). The size of the CMonly microtissues did not change by day 7, however, there was a decrease in cross sectional area observed in the CMEpiC group. Of note, the CMonly group increased circularity between day 4 and day 7 whereas the CMEpiC group demonstrated a decrease in circularity, suggesting some cellular rearrangement over the course of culture. At the day 7 endpoint, CMonly microtissues were overall more circular than the CMEpiC microtissues.

**Figure 6.**
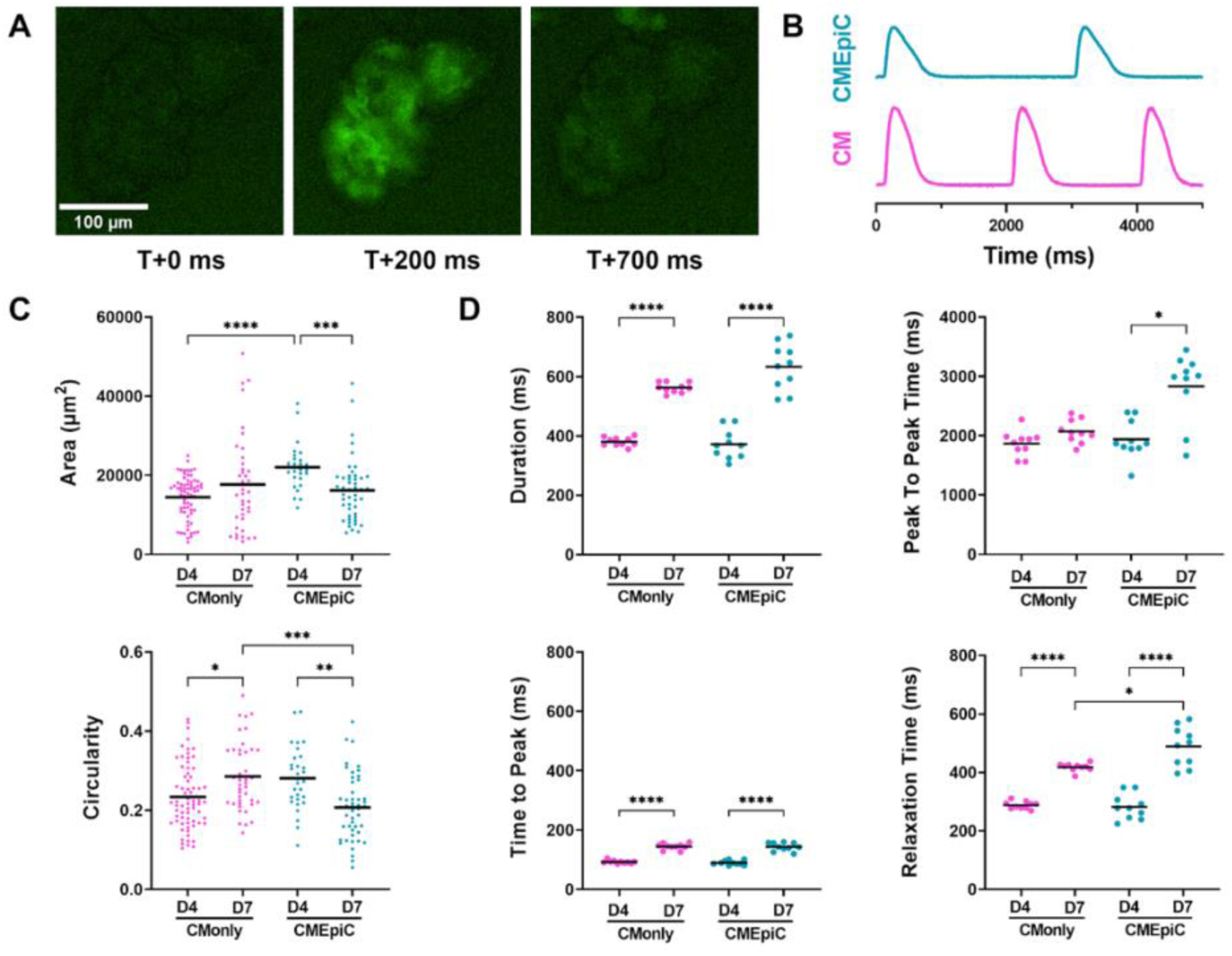
Assessment of microtissue calcium handling and morphometrics. **(A)** Time course showing the calcium flux in D7 CMEpiC microtissue. **(B)** Contraction profile comparison of D7 CMonly and D7 CMEpiC microtissues. **(C)** Area and circularity of microtissues. **(D)** Calcium flux characteristics of CMonly and CMEpiC microtissues at D4 and D7. Asterisks indicate statistically significant difference in means assessed by Brown-Forsythe and Welch ANOVA and Kruskal Wallis test: (*) p < 0.05, (**) p < 0.005, (***) p < 0.0005, (****) p < 0.0001.

Calcium handling kinetics were used as a measure of functional differences between microtissues with and without EpiC cells. When beating regions of microtissues were analyzed (Figure 6A-B), no differences were observed by day 4 of culture in contraction duration, peak-to-peak time, time to peak and relaxation time regardless of the incorporation of EpiC (Figure 6D). However, by day 7, the contraction profile had broadened in both the CMOnly and CMEpiC microtissues, with a longer peak duration, time to peak, and relaxation time. The CMEpiC microtissues also displayed a longer peak-to-peak time and relaxation time compared to the CMOnly group at D7. Overall, the addition of EpiC in this configuration produced microtissues which behaved similarly to microtissues composed only of cardiomyocytes, with EpiC not impairing the function of cardiomyocytes but promoting greater structural compaction.

## Discussion

During construction of engineered cardiac models it is important to consider all relevant cell types and investigate the role they each play in the development and maintenance of a healthy heart as well as what goes wrong during cardiac disease. Recent emphasis on the key role of the epicardium [21, 34, 36, 39, 44] has highlighted the need for more studies to examine the interplay of these cells with cardiomyocytes. In fact, the increase in labs interested in incorporating this population of cardiac cells has highlighted a need to fully understand the morphological and phenotypic changes these cells go through during in vitro culture as to ensure robust cardiac models. This study demonstrates the ease of generating large populations of epicardial cells from hiPSCs for the use of manufacturing engineered cardiac tissues. Further, key morphologic and structural changes associated with the cultured epicardial cells are identified and lead to recommendations on best use of these cells in engineered tissues.

Derivation of epicardial cells is similar to that of cardiomyocytes, but includes a critical extra step of Wnt reactivation. Previous studies have noted that a failure to reactivate Wnt in iPSC-derived cardiac progenitor cells results in sheets of beating cardiomyocytes along the lines of typical cardiomyocyte differentiation strategies [9]. Others have likewise noted that epicardial specification by coupled Wnt and BMP4 activation does not entirely suppress the cardiomyocyte-leaning subset of the cardiac progenitor population [45, 46]. Therefore, it is not surprising that the day 11 EpiC population examined in this study contained a small percentage of cardiomyocytes (Figure 1). It is likely that the D6 CPC population is nonuniform in its response to Wnt, with some individual cells or clusters requiring more GSK3 suppression to specify the WT1^+^ lineage. Overall we conclude that our EpiC differentiation produced a heterogeneous population at D11 which was primarily epicardial or epicardial-derived, expressing WT1, but which still contained a small percentage of cardiomyocytes.

Another notable mention is that previously published reports have used Accutase [9, 23] or Trypsin-like enzymes [21, 22, 34, 45–47] to dissociate *in vitro* epicardial cells. In the experiments outlined in this study, Accutase and Trypsin proved ineffective for singularization of EpiC at D11. Accutase alone dissociated the cells from the surface of the plate but failed to break the cell-cell junctions, leaving sheets of cells; Trypsin broke cell-cell junctions but did not dissociate the cells from the surface, leaving them attached to the plates. A 2:1 Accutase:Trypsin mixture achieved success, fully dissociating the cells, but required incubation times in excess of 20 minutes and neutralization in culture media containing serum. Substituting AccuMax at this and subsequent stages of EpiC differentiation improved dissociation, requiring only five minutes of incubation to achieve complete singularization and allowing neutralization with DPBS. Our recommendation is therefore to use AccuMax for rapid and efficient dissociation of iPSC-EpiC.

Although previous reports have suggested the differentiated epicardial phenotype is capable of persisting for long culture durations [9, 20], we have identified morphological size increases and limited cell expansion over the course of culture, despite maintenance of an epicardial WT1+ phenotype. This decrease in cell yield appears related to an increase in cell size, as increasingly large cells fill a culture dish of a fixed size with fewer total cells. This effect is similar to the behavior observed in senescent late-passage cultured primary cells [48]. Nevertheless, higher-passage EpiC remain robustly WT1^+^ and preserve their epithelial morphology. While it was initially hypothesized that P1 EpiC might represent a purer epicardial population, this is not the case; the proportion of WT1^+^ cells within all EpiC populations from D11 through P5 remained consistently high (Figure 2). Both D11 and P1 EpiC can be formed into spherical MT, though D11 MT were more consistent in shape over time (Figure 3). Taken alongside the decline in cell numbers over time, we conclude that for biomanufacturing purposes D11 EpiC are preferable: D11 EpiC require less time and volume of reagents to produce compared to later-passage cells, more EpiC can be obtained at D11 than at subsequent passages for a given growth area, D11 EpiC have an equally high WT1+ percentage, and D11 EpiC form into MT more robustly than P1 EpiC. Therefore, low-passage EpiC cells represent a population that more closely mimics the cells found in the native epicardium.

During cardiac development cells from the proepicardial organ form the epicardium by migration over the surface of the primitive myocardium [4]. iPSC-EpiC assembled into 3D MT preserve this capacity, expanding across a 2D surface and forming a monolayer.

These cells maintained their phenotype as indicated by robust WT1 expression in the migrating population. The migrating EpiC also displayed an increase in size, consistent with murine and zebrafish epicardial regeneration [8]. A similar regenerative mechanism has also been reported in *Drosophila* epithelium and murine endothelium [8, 49, 50]. The presence of multinucleated cells within EpiC monolayers (apparent in Figure 5A) is consistent with the ploidy-driven regeneration mechanism described in the animal models. Interestingly, the population nearest to the expansion wavefront displayed a greater proportion of large cells compared to the population nearest to the MT. The existence of these “Leader” and “Follower” subpopulations indicates that the migratory behavior reported in zebrafish and mouse epicardium could be preserved in human EpiC [8]. The size difference between the two subpopulations could also account for the variation in cell size observed in 2D EpiC culture. The presence of more “Leader” EpiC in a given population would produce a greater mean cell size, and in the immunofluorescence images in this study a variety of cell size and shape among EpiC was observed, supporting this hypothesis. It is also known that epicardial cells can become activated and differentiate into multiple stromal populations in the heart including fibroblasts and smooth muscle cells [25–27, 44]. It is possible that the dual population observed here with the “Leaders” and “Followers” may give rise to different stromal cells based on the chemical and physical cues they receive.

Since the heart is a complex microenvironment including multiple extracellular matrix molecules [1], it was important to determine if ECM coating played a role in the maintenance of EpiC phenotype. In fact, the difference in EpiC migration over various extracellular matrix coatings was substantial. By day 3, all samples exhibited a similar degree of circularity (overall mean 0.82) indicating a broadly uniform expansion however, surfaces treated with 5 μg/cm^2^ laminin produced the largest skirt area, indicating a migratory advantage. Laminin is one of the ECM molecules that is most abundant in the heart [51] and therefore and may play a role in supporting EpiC migration in vivo. TCPS consistently ranked last in mean skirt area throughout the culture, indicating that without an ECM coating, EpiC migration and outgrowth is stunted. These results are consistent with the general behavior of EpiC, reflecting a cell type which can readily migrate within an extracellular matrix environment or produce its own [21, 52, 53, 54, 55]. For biomanufacturing purposes, 10 µg/cm^2^ gelatin is the most cost-effective option for large scale culture of EpiC, and was found to consistently outperform TCPS. 5 μg/cm^2^ laminin was found to produce the largest skirt, and may be preferable to gelatin for applications where a large number of EpiC are required quickly.

To ensure that the EpiC derived in this study were amenable to fabricating 3D cardiac tissues, spherical aggregates were formed, analogous to many other cardiac tissue engineering studies [21, 34–36, 39]. The addition of EpiC caused no impedance to the function of these cardiac tissues at early time points. Previous studies have constructed similar spherical microtissues using iPSC-derived EpiC, forming complete co-cultured structures between 14 and 21 days of culture [34, 35] and these tissues have required 14 days of culture to reach a stable structural state even with a favorable extracellular matrix environment [21]. This suggests that longer culture may be necessary to realize the full benefit of epicardial cell incorporation into complex engineered tissues.

## Conclusions

In this study, we explore the use of iPSC-derived epicardial populations for studying heart development and fabricating heterotypic engineered tissue constructs. Further, we identify specific recommendations for the differentiation and maintenance of epicardial populations, including use of low-passage cells, use of Accutase for passaging, and culture on laminin-coated surfaces. Taken together, these recommendations lead to a robust population of epicardial cells that can be easily incorporated into 3D engineered cardiac tissues.

## Acknowledgements

The authors would like to acknowledge Natalie Pachter and Muhammad Arslan Tayyab for technical assistance on experimental technique; Dr. Bruce Conklin, who generously provided the GCaMP6f-modified iPSC cell line; and Dr. Yan Sun and the staff of the Binghamton University Health Science Core Facility and Advanced Diagnostics Laboratory. Schematics were produced using BioRender. Financial support for this research was provided by the National Institutes of Health (National Heart, Lung, and Blood Institute Grant # 1R15HL140745) and by Binghamton University.

## Author Contributions

K.B., S.A., and T.A.H. designed the experiments; K.B. and S.A. performed the experiments and completed data processing; J.J. and K.B. wrote the J-SEG MATLAB code; K.B, S.A., and T.A.H. wrote and edited the manuscript.

## Ethics Statement

All pluripotent stem cell lines used are publicly available and their derivation protocols are documented by the suppliers.

## Data Availability Statement

All raw data from this work is available upon request of the authors.

---

**Supplementary Table 1:**
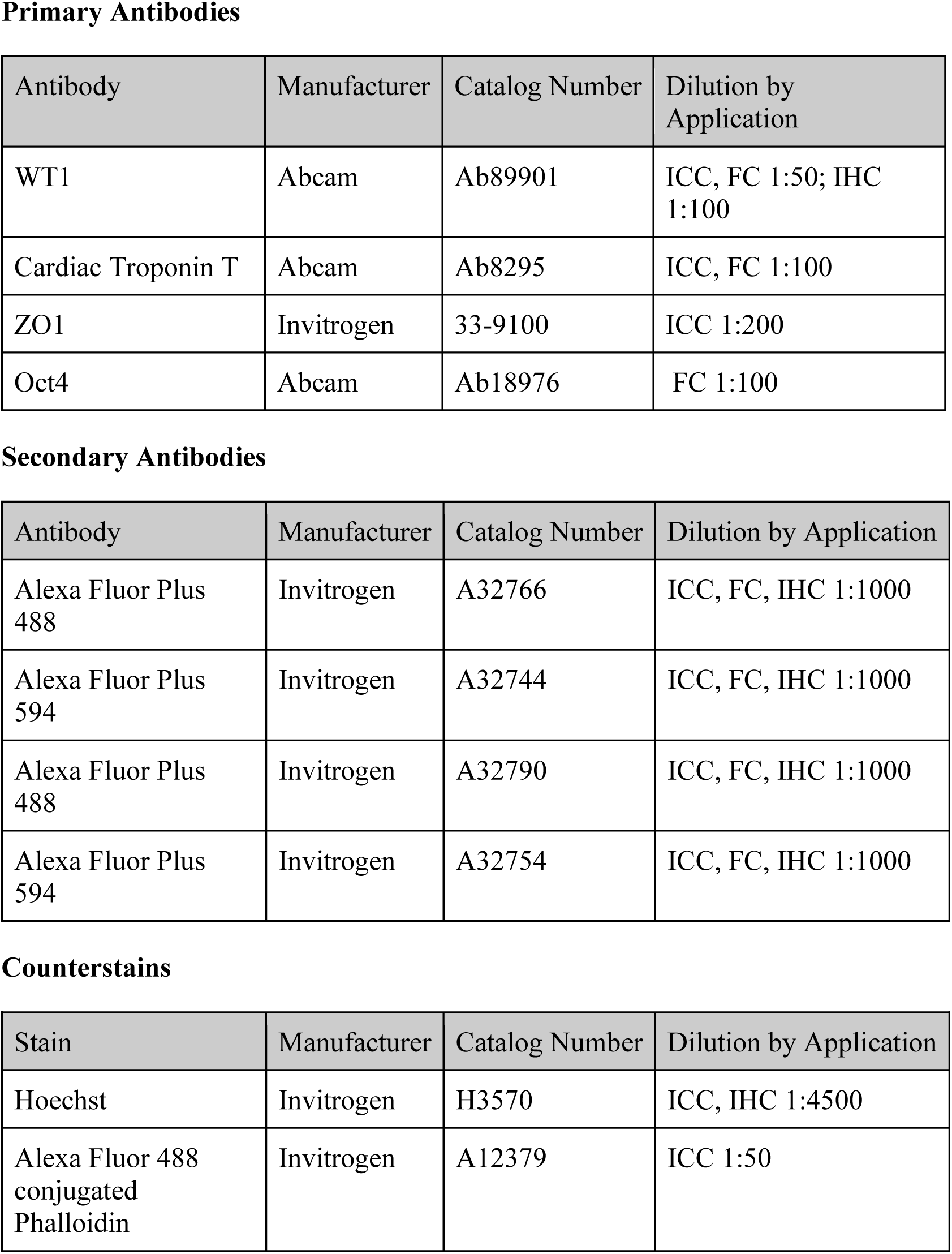
Immunostaining Reagents.

**Supplementary Table 2:**
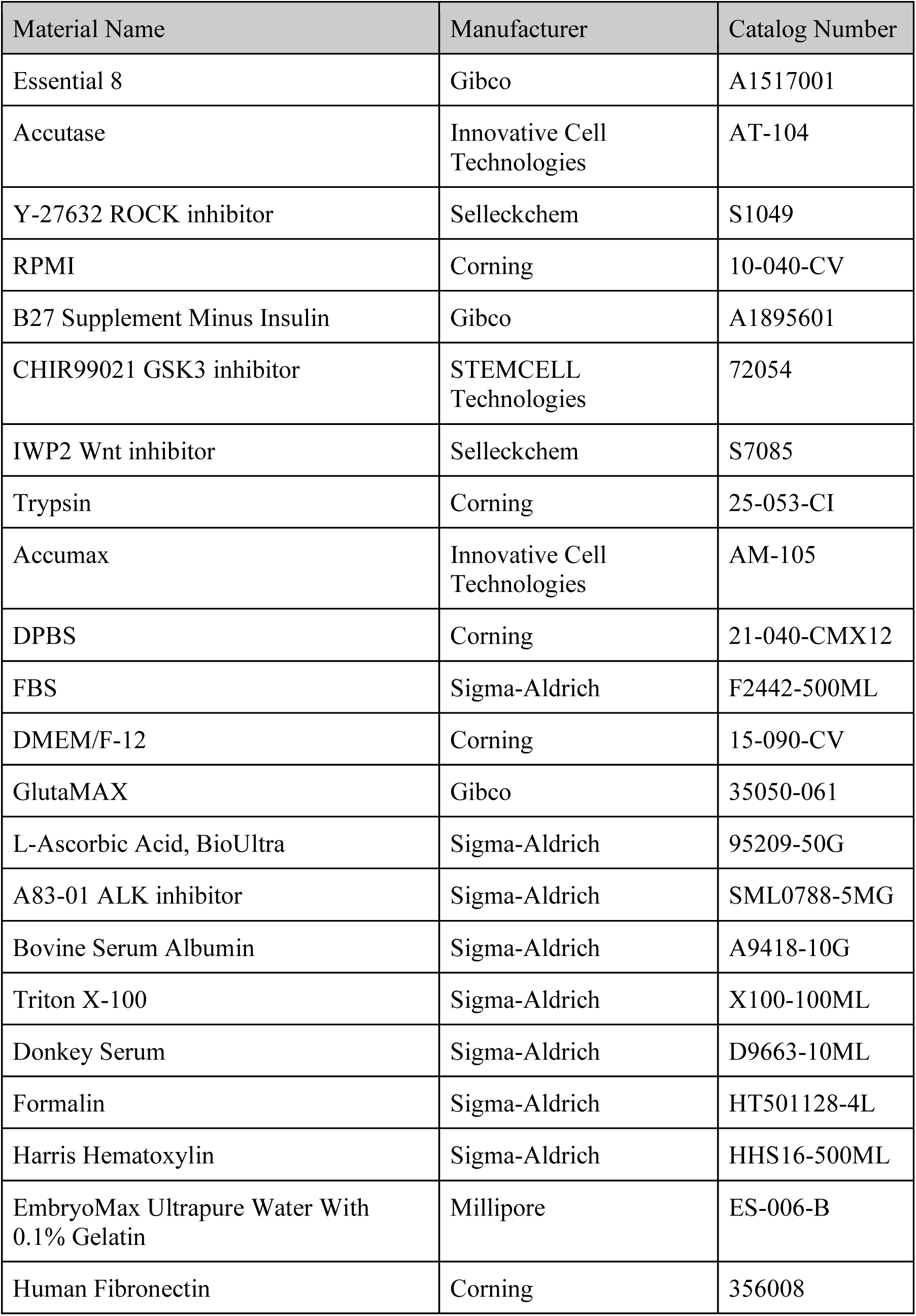
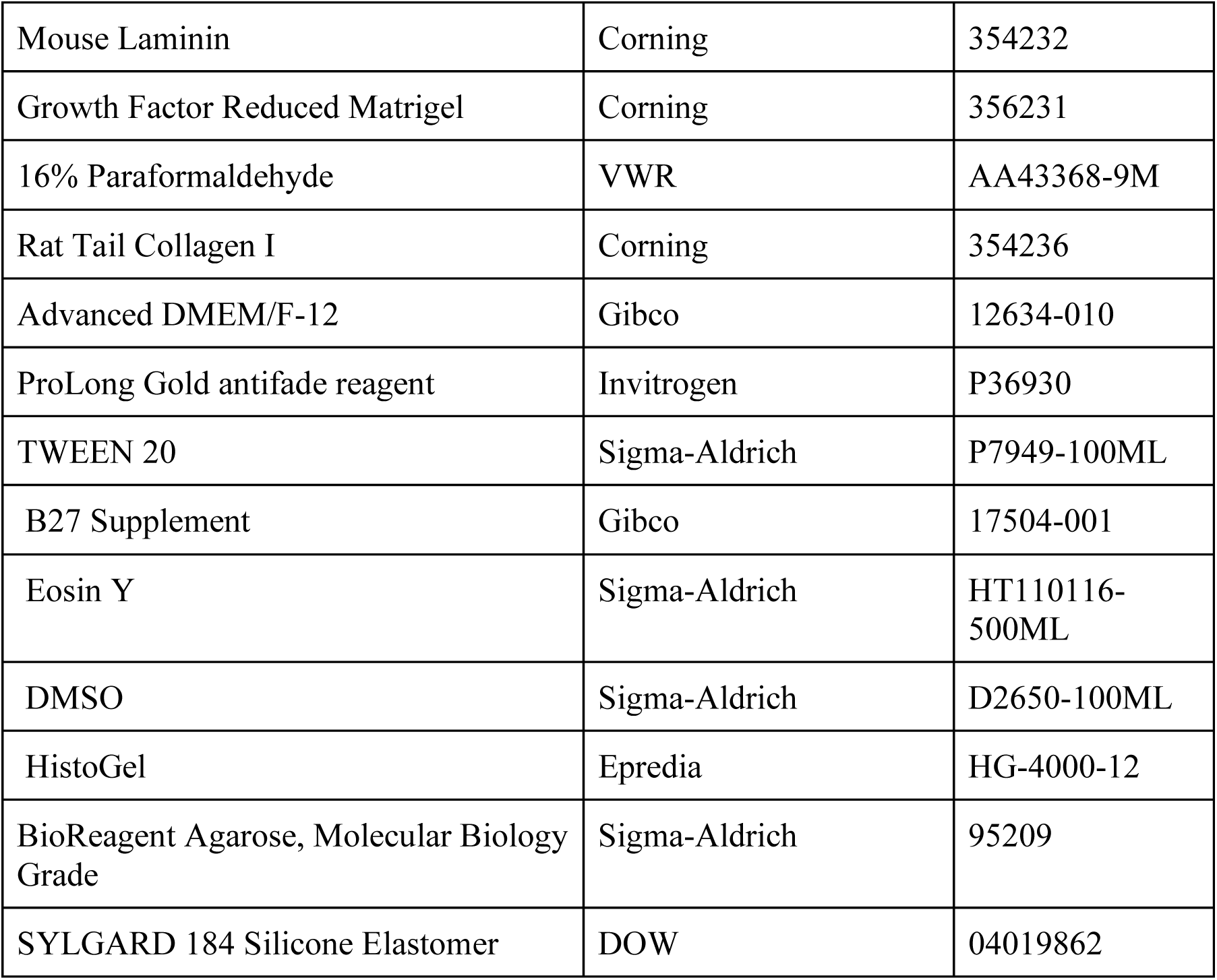
Reagent Information.

## References

[1] A. C. Silva, C. Pereira, A. C. R. G. Fonseca, P. Pinto-do-Ó, and D. S. Nascimento, “Bearing My Heart: The Role of Extracellular Matrix on Cardiac Development, Homeostasis, and Injury Response,” Front. Cell Dev. Biol., vol. 8, p. 621644, Jan. 2021, doi: 10.3389/fcell.2020.621644.

[2] X. Fu et al., “Specialized fibroblast differentiated states underlie scar formation in the infarcted mouse heart,” J Clin Invest, vol. 128, no. 5, pp. 2127–2143, May 2018, doi: 10.1172/JCI98215.

[3] M. D. Tallquist and J. D. Molkentin, “Redefining the identity of cardiac fibroblasts,” Nat Rev Cardiol, vol. 14, no. 8, pp. 484–491, Aug. 2017, doi: 10.1038/nrcardio.2017.57.

[4] L. S. Rodgers, S. Lalani, R. B. Runyan, and T. D. Camenisch, “Differential growth and multicellular villi direct proepicardial translocation to the developing mouse heart,” Developmental Dynamics, vol. 237, no. 1, pp. 145–152, 2008, doi: 10.1002/dvdy.21378.

[5] J. Schlueter and T. Brand, “Epicardial Progenitor Cells in Cardiac Development and Regeneration,” J. of Cardiovasc. Trans. Res., vol. 5, no. 5, pp. 641–653, Oct. 2012, doi: 10.1007/s12265-012-9377-4.

[6] P. C. Nahirney, T. Mikawa, and D. A. Fischman, “Evidence for an extracellular matrix bridge guiding proepicardial cell migration to the myocardium of chick embryos,” Developmental Dynamics, vol. 227, no. 4, pp. 511–523, Aug. 2003, doi: 10.1002/dvdy.10335.

[7] C. A. Risebro, J. M. Vieira, and P. R. Riley, “Characterisation of the human embryonic and foetal epicardium during heart development,” *Development*, p. dev.127621, Jan. 2015, doi: 10.1242/dev.127621.

[8] J. Cao et al., “Tension Creates an Endoreplication Wavefront that Leads Regeneration of Epicardial Tissue,” Dev Cell, vol. 42, no. 6, pp. 600–615.e4, Sep. 2017, doi: 10.1016/j.devcel.2017.08.024.

[9] X. Bao et al., “Long-term self-renewing human epicardial cells generated from pluripotent stem cells under defined xeno-free conditions,” Nature Biomedical Engineering, vol. 1, no. 1, Art. no. 1, Dec. 2016, doi: 10.1038/s41551-016-0003.

[10] A. W. Moore, L. McInnes, J. Kreidberg, N. D. Hastie, and A. Schedl, “YAC complementation shows a requirement for *Wt1* in the development of epicardium, adrenal gland and throughout nephrogenesis,” Development, vol. 126, no. 9, pp. 1845–1857, May 1999, doi: 10.1242/dev.126.9.1845.

[11] S. N. Duim, A. M. Smits, B. P. T. Kruithof, and M.-J. Goumans, “The roadmap of WT1 protein expression in the human fetal heart,” Journal of Molecular and Cellular Cardiology, vol. 90, pp. 139–145, Jan. 2016, doi: 10.1016/j.yjmcc.2015.12.008.

[12] X. Lian et al., “Robust cardiomyocyte differentiation from human pluripotent stem cells via temporal modulation of canonical Wnt signaling,” Proc Natl Acad Sci U S A, vol. 109, no. 27, pp. E1848–1857, Jul. 2012, doi: 10.1073/pnas.1200250109.

[13] P. W. Burridge, G. Keller, J. D. Gold, and J. C. Wu, “Production of De Novo Cardiomyocytes: Human Pluripotent Stem Cell Differentiation and Direct Reprogramming,” Cell Stem Cell, vol. 10, no. 1, pp. 16–28, Jan. 2012, doi: 10.1016/j.stem.2011.12.013.

[14] G. H. Hwang et al., “Purification of small molecule-induced cardiomyocytes from human induced pluripotent stem cells using a reporter system,” Journal of cellular physiology, vol. 232, no. 12, pp. 3384–3395, Dec. 2017, doi: 10.1002/jcp.25783.

[15] S. J. Kattman, T. L. Huber, and G. M. Keller, “Multipotent Flk-1+ Cardiovascular Progenitor Cells Give Rise to the Cardiomyocyte, Endothelial, and Vascular Smooth Muscle Lineages,” Developmental Cell, vol. 11, no. 5, pp. 723–732, Nov. 2006, doi: 10.1016/j.devcel.2006.10.002.

[16] J. H. Lee, S. I. Protze, Z. Laksman, P. H. Backx, and G. M. Keller, “Human Pluripotent Stem Cell-Derived Atrial and Ventricular Cardiomyocytes Develop from Distinct Mesoderm Populations,” Cell Stem Cell, vol. 21, no. 2, pp. 179–194.e4, Aug. 2017, doi: 10.1016/j.stem.2017.07.003.

[17] V. Schwach and R. Passier, “Generation and purification of human stem cell-derived cardiomyocytes,” Differentiation, vol. 91, no. 4, pp. 126–138, Apr. 2016, doi: 10.1016/j.diff.2016.01.001.

[18] C. Xu, S. Police, N. Rao, and M. K. Carpenter, “Characterization and Enrichment of Cardiomyocytes Derived From Human Embryonic Stem Cells,” Circulation Research, vol. 91, no. 6, pp. 501–508, Sep. 2002, doi: 10.1161/01.RES.0000035254.80718.91.

[19] G. E. Brown et al., “Engineered cocultures of iPSC-derived atrial cardiomyocytes and atrial fibroblasts for modeling atrial fibrillation,” Sci. Adv., vol. 10, no. 3, p. eadg1222, Jan. 2024, doi: 10.1126/sciadv.adg1222.

[20] X. Bao, X. Lian, T. Qian, V. J. Bhute, T. Han, and S. P. Palecek, “Directed differentiation and long-term maintenance of epicardial cells from human pluripotent stem cells under fully defined conditions,” Nat Protoc, vol. 12, no. 9, pp. 1890–1900, Sep. 2017, doi: 10.1038/nprot.2017.080.

[21] J. Bargehr et al., “Epicardial cells derived from human embryonic stem cells augment cardiomyocyte-driven heart regeneration,” Nat Biotechnol, vol. 37, no. 8, pp. 895–906, Aug. 2019, doi: 10.1038/s41587-019-0197-9.

[22] D. Iyer et al., “Robust derivation of epicardium and its differentiated smooth muscle cell progeny from human pluripotent stem cells,” Development, vol. 142, no. 8, pp. 1528–1541, Apr. 2015, doi: 10.1242/dev.119271.

[23] X. Bao, V. J. Bhute, T. Han, T. Qian, X. Lian, and S. P. Palecek, “Human pluripotent stem cell-derived epicardial progenitors can differentiate to endocardial-like endothelial cells,” Bioengineering & Translational Medicine, vol. 2, no. 2, pp. 191– 2011, 2017, doi: 10.1002/btm2.10062.

[24] C. Z. Liu et al., “Feeder-free generation and characterization of endocardial and cardiac valve cells from human pluripotent stem cells,” iScience, vol. 27, no. 1, p. 108599, Jan. 2024, doi: 10.1016/j.isci.2023.108599.

[25] A. J. Whitehead, J. D. Hocker, B. Ren, and A. J. Engler, “Improved epicardial cardiac fibroblast generation from iPSCs,” Journal of Molecular and Cellular Cardiology, vol. 164, pp. 58–68, Mar. 2022, doi: 10.1016/j.yjmcc.2021.11.011.

[26] M. E. Floy et al., “Developmental lineage of human pluripotent stem cell-derived cardiac fibroblasts affects their functional phenotype,” The FASEB Journal, vol. 35, no. 9, p. e21799, Sep. 2021, doi: 10.1096/fj.202100523R.

[27] F. T. Bekedam et al., “Mechanical stimulation of induced pluripotent stem derived cardiac fibroblasts,” Sci Rep, vol. 14, no. 1, p. 9795, Apr. 2024, doi: 10.1038/s41598-024-60102-w.

[28] S. I. Protze et al., “Sinoatrial node cardiomyocytes derived from human pluripotent cells function as a biological pacemaker,” Nat Biotechnol, vol. 35, no. 1, pp. 56–68, Jan. 2017, doi: 10.1038/nbt.3745.

[29] L. Yin et al., “RA signaling pathway combined with Wnt signaling pathway regulates human-induced pluripotent stem cells (hiPSCs) differentiation to sinus node-like cells,” Stem Cell Res Ther, vol. 13, no. 1, p. 324, Dec. 2022, doi: 10.1186/s13287-022-03006-8.

[30] H. Zhao et al., “Overexpression of TBX3 in human induced pluripotent stem cells (hiPSCs) increases their differentiation into cardiac pacemaker-like cells,” Biomedicine & Pharmacotherapy, vol. 130, p. 110612, Oct. 2020, doi: 10.1016/j.biopha.2020.110612.

[31] E. Giacomelli et al., “Human-iPSC-Derived Cardiac Stromal Cells Enhance Maturation in 3D Cardiac Microtissues and Reveal Non-cardiomyocyte Contributions to Heart Disease,” Cell Stem Cell, vol. 26, no. 6, pp. 862–879.e11, Jun. 2020, doi: 10.1016/j.stem.2020.05.004.

[32] D. C. Nguyen et al., “Microscale Generation of Cardiospheres Promotes Robust Enrichment of Cardiomyocytes Derived from Human Pluripotent Stem Cells,” Stem Cell Reports, vol. 3, no. 2, pp. 260–268, Aug. 2014, doi: 10.1016/j.stemcr.2014.06.002.

[33] T. A. Hookway, O. B. Matthys, D. A. Joy, J. E. Sepulveda, R. Thomas, and T. C. McDevitt, “Bi-directional Impacts of Heterotypic Interactions in Engineered 3D Human Cardiac Microtissues Revealed by Single-Cell RNA-Sequencing and Functional Analysis.” Jul. 06, 2020. doi: 10.1101/2020.07.06.190504.

[34] J. J. Tan et al., “Human iPS-derived pre-epicardial cells direct cardiomyocyte aggregation expansion and organization in vitro,” Nat Commun, vol. 12, no. 1, p. 4997, Aug. 2021, doi: 10.1038/s41467-021-24921-z.

[35] E. Giacomelli et al., “Human-iPSC-Derived Cardiac Stromal Cells Enhance Maturation in 3D Cardiac Microtissues and Reveal Non-cardiomyocyte Contributions to Heart Disease,” Cell Stem Cell, vol. 26, no. 6, pp. 862–879.e11, Jun. 2020, doi: 10.1016/j.stem.2020.05.004.

[36] A. C. Silva et al., “Developmental co-emergence of cardiac and gut tissues modeled by human iPSC-derived organoids,” Developmental Biology, preprint, May 2020. doi: 10.1101/2020.04.30.071472.

[37] C. V. Del Campo et al., “Regenerative potential of epicardium-derived extracellular vesicles mediated by conserved miRNA transfer,” Cardiovascular Research, vol. 118, no. 2, pp. 597–611, Jan. 2022, doi: 10.1093/cvr/cvab054.

[38] D. Bannerman et al., “Heart-on-a-Chip Model of Epicardial-Myocardial Interaction in Ischemia Reperfusion Injuryz,” Adv Healthcare Materials, p. 2302642, Apr. 2024, doi: 10.1002/adhm.202302642.

[39] L. P. Ong et al., “Epicardially secreted fibronectin drives cardiomyocyte maturation in 3D-engineered heart tissues,” Stem Cell Reports, vol. 18, no. 4, pp. 936–951, Apr. 2023, doi: 10.1016/j.stemcr.2023.03.002.

[40] T. A. Hookway et al., “Phenotypic Variation Between Stromal Cells Differentially Impacts Engineered Cardiac Tissue Function,” Tissue Eng Part A, vol. 25, no. 9–10, pp. 773–785, May 2019, doi: 10.1089/ten.tea.2018.0362.

[41] E. Hodneland, T. Kögel, D. M. Frei, H.-H. Gerdes, and A. Lundervold, “CellSegm - a MATLAB toolbox for high-throughput 3D cell segmentation,” Source Code Biol Med, vol. 8, no. 1, p. 16, Dec. 2013, doi: 10.1186/1751-0473-8-16.

[42] T. A. Hookway, J. C. Butts, E. Lee, H. Tang, and T. C. McDevitt, “Aggregate formation and suspension culture of human pluripotent stem cells and differentiated progeny,” Methods, vol. 101, pp. 11–20, May 2016, doi: 10.1016/j.ymeth.2015.11.027.

[43] L. Sala et al., “MUSCLEMOTION: A Versatile Open Software Tool to Quantify Cardiomyocyte and Cardiac Muscle Contraction In Vitro and In Vivo,” Circulation Research, vol. 122, no. 3, Feb. 2018, doi: 10.1161/CIRCRESAHA.117.312067.

[44] I. Fernandes, S. Funakoshi, H. Hamidzada, S. Epelman, and G. Keller, “Modeling cardiac fibroblast heterogeneity from human pluripotent stem cell-derived epicardial cells,” Nat Commun, vol. 14, no. 1, p. 8183, Dec. 2023, doi: 10.1038/s41467-023-43312-0.

[45] A. D. Witty et al., “Generation of the epicardial lineage from human pluripotent stem cells,” Nat Biotechnol, vol. 32, no. 10, pp. 1026–1035, Oct. 2014, doi: 10.1038/nbt.3002.

[46] J. A. Guadix et al., “Human Pluripotent Stem Cell Differentiation into Functional Epicardial Progenitor Cells,” Stem Cell Reports, vol. 9, no. 6, pp. 1754–1764, Dec. 2017, doi: 10.1016/j.stemcr.2017.10.023.

[47] H. Eid et al., “Role of epicardial mesothelial cells in the modification of phenotype and function of adult rat ventricular myocytes in primary coculture.,” Circ Res, vol. 71, no. 1, pp. 40–50, Jul. 1992, doi: 10.1161/01.RES.71.1.40.

[48] F. Rodier and J. Campisi, “Four faces of cellular senescence,” Journal of Cell Biology, vol. 192, no. 4, pp. 547–556, Feb. 2011, doi: 10.1083/jcb.201009094.

[49] V. P. Losick, D. T. Fox, and A. C. Spradling, “Polyploidization and Cell Fusion Contribute to Wound Healing in the Adult Drosophila Epithelium,” Current Biology, vol. 23, no. 22, pp. 2224–2232, Nov. 2013, doi: 10.1016/j.cub.2013.09.029.

[50] V. P. Losick, A. S. Jun, and A. C. Spradling, “Wound-Induced Polyploidization: Regulation by Hippo and JNK Signaling and Conservation in Mammals,” PLoS ONE, vol. 11, no. 3, p. e0151251, Mar. 2016, doi: 10.1371/journal.pone.0151251.

[51] Q. Jallerat and A. W. Feinberg, “Extracellular Matrix Structure and Composition in the Early Four-Chambered Embryonic Heart,” Cells, vol. 9, no. 2, p. 285, Jan. 2020, doi: 10.3390/cells9020285.

[52] F. C. Simões and P. R. Riley, “The ontogeny, activation and function of the epicardium during heart development and regeneration,” Development, vol. 145, no. 7, p. dev155994, Apr. 2018, doi: 10.1242/dev.155994.

[53] A. C. Gittenberger-de Groot, E. M. Winter, M. M. Bartelings, M. Jose Goumans, M. C. DeRuiter, and R. E. Poelmann, “The arterial and cardiac epicardium in development, disease and repair,” Differentiation, vol. 84, no. 1, pp. 41–53, Jul. 2012, doi: 10.1016/j.diff.2012.05.002.

[54] P. Quijada, M. A. Trembley, and E. M. Small, “The Role of the Epicardium During Heart Development and Repair,” Circ Res, vol. 126, no. 3, pp. 377–394, Jan. 2020, doi: 10.1161/CIRCRESAHA.119.315857.

[55] T. J. Streef and A. M. Smits, “Epicardial Contribution to the Developing and Injured Heart: Exploring the Cellular Composition of the Epicardium,” Front. Cardiovasc. Med., vol. 8, p. 750243, Sep. 2021, doi: 10.3389/fcvm.2021.750243.

