## Supplemental Table 1 for "Engineered Cardiac Microtissue Biomanufacturing Using Human Induced Pluripotent Stem Cell Derived Epicardial Cells"

**Supplementary Table 1: Immunostaining Reagents**

**Primary Antibodies**

| Antibody | Manufacturer | Catalog Number | Dilution by Application |
|---|---|---|---|
| WT1 | Abcam | Ab89901 | ICC, FC 1:50; IHC 1:100 |
| Cardiac Troponin T | Abcam | Ab8295 | ICC, FC 1:100 |
| ZO1 | Invitrogen | 33-9100 | ICC 1:200 |
| Oct4 | Abcam | Ab18976 | FC 1:100 |

**Secondary Antibodies**

| Antibody | Manufacturer | Catalog Number | Dilution by Application |
|---|---|---|---|
| Alexa Fluor Plus 488 | Invitrogen | A32766 | ICC, FC, IHC 1:1000 |
| Alexa Fluor Plus 594 | Invitrogen | A32744 | ICC, FC, IHC 1:1000 |
| Alexa Fluor Plus 488 | Invitrogen | A32790 | ICC, FC, IHC 1:1000 |
| Alexa Fluor Plus 594 | Invitrogen | A32754 | ICC, FC, IHC 1:1000 |

**Counterstains**

| Stain | Manufacturer | Catalog Number | Dilution by Application |
|---|---|---|---|
| Hoechst | Invitrogen | H3570 | ICC, IHC 1:4500 |
| Alexa Fluor 488 conjugated Phalloidin | Invitrogen | A12379 | ICC 1:50 |
