## Supplemental Table 2 for "Engineered Cardiac Microtissue Biomanufacturing Using Human Induced Pluripotent Stem Cell Derived Epicardial Cells"

**Supplementary Table 2: Reagent Information**

| Material Name | Manufacturer | Catalog Number |
|---|---|---|
| Essential 8 | Gibco | A1517001 |
| Accutase | Innovative Cell Technologies | AT-104 |
| Y-27632 ROCK inhibitor | Selleckchem | S1049 |
| RPMI | Corning | 10-040-CV |
| B27 Supplement Minus Insulin | Gibco | A1895601 |
| CHIR99021 GSK3 inhibitor | STEMCELL Technologies | 72054 |
| IWP2 Wnt inhibitor | Selleckchem | S7085 |
| Trypsin | Corning | 25-053-CI |
| Accumax | Innovative Cell Technologies | AM-105 |
| DPBS | Corning | 21-040-CMX12 |
| FBS | Sigma-Aldrich | F2442-500ML |
| DMEM/F-12 | Corning | 15-090-CV |
| GlutaMAX | Gibco | 35050-061 |
| L-Ascorbic Acid, BioUltra | Sigma-Aldrich | 95209-50G |
| A83-01 ALK inhibitor | Sigma-Aldrich | SML0788-5MG |
| Bovine Serum Albumin | Sigma-Aldrich | A9418-10G |
| Triton X-100 | Sigma-Aldrich | X100-100ML |
| Donkey Serum | Sigma-Aldrich | D9663-10ML |
| Formalin | Sigma-Aldrich | HT501128-4L |
| Harris Hematoxylin | Sigma-Aldrich | HHS16-500ML |
| EmbryoMax Ultrapure Water With 0.1% Gelatin | Millipore | ES-006-B |
| Human Fibronectin | Corning | 356008 |

| | | |
|---|---|---|
| Mouse Laminin | Corning | 354232 |
| Growth Factor Reduced Matrigel | Corning | 356231 |
| 16% Paraformaldehyde | VWR | AA43368-9M |
| Rat Tail Collagen I | Corning | 354236 |
| Advanced DMEM/F-12 | Gibco | 12634-010 |
| ProLong Gold antifade reagent | Invitrogen | P36930 |
| TWEEN 20 | Sigma-Aldrich | P7949-100ML |
| B27 Supplement | Gibco | 17504-001 |
| Eosin Y | Sigma-Aldrich | HT110116-500ML |
| DMSO | Sigma-Aldrich | D2650-100ML |
| HistoGel | Epredia | HG-4000-12 |
| BioReagent Agarose, Molecular Biology Grade | Sigma-Aldrich | 95209 |
| SYLGARD 184 Silicone Elastomer | DOW | 04019862 |
