## Supplemental Table 3 for "Engineered Cardiac Microtissue Biomanufacturing Using Human Induced Pluripotent Stem Cell Derived Epicardial Cells"

**Abbreviations**

| Abbreviation | Long Form |
|---|---|
| PEO | Proepicardial Organ |
| MT | Microtissue |
| ECM | Extracellular Matrix |
| TCPS | Tissue Culture Polystyrene |
| EpiC | Epicardial |
| CM | Cardiomyocyte |
| CF | Cardiac Fibroblast |
| fCF | Fetal Cardiac Fibroblast |
| hCF | Human Cardiac Fibroblast |
| CMonly | Cardiomyocyte Only |
| CMEpiC | Cardiomyocyte and Epicardial |
| CMCF | Cardiomyocyte and Cardiac Fibroblast |
| WT1 | Wilm's Tumor 1 |
| cTnT | Cardiac Troponin T |
| GSK3 | Glycogen synthase kinase 3 |
| DMSO | Dimethyl Sulfoxide |

| | |
|---|---|
| ALK | Anaplastic Lymphoma Kinase |
| EDTA | disodium ethylenediaminetetraacetic acid |
| Oct4 | Octamer-binding transcription factor 4 |
